# Medial olivocochlear efferent modulation of cochlear micromechanics requires P2X4 expression in outer hair cells

**DOI:** 10.1101/2025.05.15.654229

**Authors:** Coline Riffault, Steven Condamine, Yohan Bouleau, Eric Boué-Grabot, Didier Dulon

**Affiliations:** University of Bordeaux, CNRS, IMN, UMR 5293, F-33000 Bordeaux, France; Institut de l’Audition, Inserm UA1335, Centre de Recherche et d’Innovation en Audiologie Humaine (CERIAH), Institut Pasteur, F-75015 Paris, France; Bordeaux Neurocampus, University of Bordeaux, F-33000 Bordeaux, France

**Author notes:** Should be considered as co-lead authors (Eric Boué-Grabot, Didier Dulon). **Correspondence should be addressed to:** Didier Dulon, Éric Boué-Grabot. **Author Contributions:** EBG and DD designed research, CR, SC and YB performed research, CR, SC and YB analyzed data, CR, EBG and DD wrote the paper.

## Abstract

The role of P2X4, one of the most abundant ionotropic purinergic receptors in the central nervous system, is explored here in the context of auditory function. We observed, by using constitutive and conditional P2X4mCherryIN knock-in adult mouse models, a specific high expression of mCherry-tagged P2X4 in living cochlear outer hair cells (OHCs), from immature postnatal stages to adulthood. This P2X4-mCherry expression, confirmed by immuno-confocal fluorescence microscopy in wild-type mice, was mainly concentrated in the intracellular apical region of the OHCs, in the area of the Hensen’s body, a lysosomal rich region, specifically labeled with the fluorescent dye lysotracker. In addition, the basal cholinergic efferent synaptic region of the OHCs was found to express P2X4 at the cell membrane. Surprisingly, the assessment of the hearing function in constitutive P2X4 knock-out (P2X4KO) mice showed improved auditory brainstem responses with smaller latencies, larger amplitudes and smaller thresholds. These P2X4KO mice, as well as conditional P2X4KO-Myo15-Cre mice, displayed enhanced distortion product otoacoustic emissions (DPOAEs), suggesting an improved electromechanical ‘amplification’ activity by OHCs. These mutant animals showed reduced inhibition of DPOAEs by contralateral noise, consistent with a weaker inhibitory effect of the medial cholinergic olivocochlear efferent circuit (MOC) on OHCs. We concluded that the MOC negative feedback modulation of cochlear micromechanics, in addition to involve Ca^2+^ permeable α9/α10 nicotinic receptors, also requires the activation of postsynaptic P2X4 receptors in OHCs.

**SIGNIFICANCE STATEMENT:** Our study reveals a specific strong expression of P2X4 in mouse cochlear outer hair cells (OHCs), these cells being essential for generating the distortion products otoacoustic emissions (DPOAES) and tuning the sensitivity and frequency selectivity of the cochlea. Mice lacking P2X4 showed a deficient inhibitory control of their cochlear DPOAEs when activating the medial olivocochlear efferent pathway innervating the OHCs. We propose P2X4 receptors as an important Ca^2+^ regulatory component of the micromechanics of OHCs and that genetic defects in these purinergic receptors may potentially lead to hearing disorders such as tinnitus and hyperacusis.

## INTRODUCTION

P2X receptors are cation-permeable ligand-gated ion channels that open upon binding of extracellular adenosine 5’-triphosphate (ATP). These purinergic receptors largely contribute to Ca^2+^ entry in excitable and non-excitable cells (Egan and Khakh, 2004). Seven genes coding for P2X subunits have been identified and named as P2X1 to P2X7. Their expression controls a wide variety of physiological process from muscle contraction to synaptic transmission in central and peripheral sensory neurons (North, 2002; Khakh and North 2012).

Of the seven P2X subunits, P2X4 receptors (P2X4) are the most abundant in the central nervous system (CNS) and are expressed in neurons across multiple regions in the brain and spinal cord as well as in microglia. In normal condition, P2X4 is constitutively internalized and as a result, P2X4 is found preferentially in intracellular compartments and poorly expressed in cell membrane surface. In contrast, in various conditions and neurological diseases such as chronic pain, anxiety, multiple sclerosis and neurodegenerative diseases, surface P2X4 receptors are upregulated by mobilization of intracellular pools and/or *de novo* expression in glia or neurons (Duveau et al., 2020; Montilla et al., 2020). The specific increase in surface P2X4 expression in spinal microglia is critical for the pathogenesis of chronic pain (Tsuda et al. 2003; Beggs et al., 2012; Ulmann et al., 2008).

In cochlear auditory physiology, the expression and function of P2X2 and P2X7 are already established in the organ of Corti, the vestibular organ and the rat spiral ganglion at different stages of postnatal development (Jovanovic and Milenkovic, 2020). In particular, these receptors have been shown to control the spontaneous electrical activity of hair cells during development, as well as their receptor potentials during sound stimulation in the adult stage (Ito and Dulon, 2010; Wang et al., 2015; Cederholm et al., 2019). P2X4 expression was recently found in mouse cochlear hair cells by RNA-seq techniques (Orvis et al., 2021). Noise exposure was shown to increase P2X4 expression in guinea pigs’ cochlear spiral ganglion neurons (Shi et al., 2023), but the precise role of these purinergic receptors in auditory function remains unknown. To address this question, we characterized the auditory function of constitutive and various conditional P2X4 knockout (P2X4KO) mouse models. Using P2X4mCherryIN knock-in (P2X4KI) mice (Bertin et al 2021), we demonstrated here a strong expression of P2X4 in outer hair cells (OHCs) of the organ of Corti. Theses purinergic receptors were here found to be essential in controlling, via the medial olivocochlear efferent system (MOC) innervating the OHCs, the distortion product otoacoustic emissions (DPOAEs). In the cochlea, DPOAEs are generated by the electromotile OHCs when two pure tones (f1, f2) are presented simultaneously to the ear (Kemp, 2002; Brownell, 1990; Dulon and Schacht 1992; Dallos, 2008; Ashmore et al., 2011). DPOAEs reflect an active cochlear process that refines the sensitivity and frequency selectivity of the mechanical vibrations of the basilar membrane supporting the organ of Corti.

## MATERIAL ET METHODS

### Mouse lines

To elucidate P2X4 functions, innovative P2X4mCherryIN knock-in mice were developed, showing an increase in the number of P2X4 receptors at the surface plasma membrane via the deletion of its endocytosis site replaced by the mCherry fluorescent protein (Bertin et al., 2021). General P2X4KI mice, i.e. P2X4mCherryIN, where wild-type (WT) P2X4 is constitutively substituted by P2X4mCherryIN in all cells expressing natively P2X4 were used. General P2X4KI mice were obtained initially by breeding floxed P2X4mCherryIN^fl/fl^ (expressing P2X4WT) with cytomegalovirus (CMV) promoter-Cre mice. The ubiquitous activity of the CMV promotor induced the excision of the floxed cassette in the germinal cell lines and direct transmission to offspring. By breeding these animals, we thus obtained a constitutive P2X4mCherryIN^Exc/Exc^ mouse line (namely P2X4KI or P2X4mCherryIN). To allow substitution of P2X4 by P2X4mCherryIN in specific cell populations, we used floxed P2X4mCherryIN^fl/fl^ and Cre activity-recombinase under the control of specific promoters: 1) Synapsin1-Cre^+/−^:P2X4mCherryIN^fl/fl^ (neuron specific; Klüger et al., 2003), and Myo15-Cre^+/−^:P2X4mCherryIN^fl/fl^ (hair cell specific, Dulon et al., 2018). Finally, we also studied general P2X4KO mice and conditional P2X4KO mice by breeding floxed P2X4^fl/fl^ mice (Gilabert et al, 2023) and Myo15-Cre^+/+^ mice to obtain P2X4KO^fl/fl^:Myo15-Cre^+/−^ mice. All the mice used during this project were maintained on a pure C57BL/6 background. All hearing tests were conducted on mice no older than P60 since the cadherin-23 mutation causes these mice to exhibit early hearing loss (Peineau et al., 2021).

### Auditory function testing: ABRs and DPOAEs

Experiments were performed in C57BL6/J mice of either sex according to the ethical guidelines of the University of Bordeaux (CE050; APAFIS#22860) and French Ministry of Agriculture agreement (C33-063-075). Measurements of Auditory Brainstem Responses (ABRs) and Distortion Products OtoAcoustic Emissions (DPOAEs) were made on isoflurane anesthetized male and female mice, aged of 30-60 days (P30, P60). To record ABRs, which represent the sound-evoked synchronous firing of the auditory cochlear nerve fibers and the activation of the subsequent central auditory relays), anesthetized mice were placed in a closed anechoic recording chamber, with its body temperature kept constant at 37°C. Sound stimulus generation and data acquisition, we performed with a TDT RZ6/BioSigRZ system coupled to an EC-1 electrostatic driver to generate primary tones (Tucker-Davis Technologies). Click or tone-based ABR signals were averaged after the sound presentation of a series of 512 stimulations. Thresholds were defined as the lowest stimulus for recognizable wave-I and II. The amplitude of ABR wave-I was estimated by measuring the voltage difference between the positive (P1) and negative (N1) peak of wave-I. Sound intensities of 10 to 90 dB SPL in 10 dB step, were tested.

DPOAEs, which originate from the electromechanical activity of the outer hair cells (OHCs), were tested by using two simultaneous continuous pure tones with frequency ratio of 1.2 (f1 = 12.73 kHz and f2 = 15,26 kHz) at tone level ranging from 30 to 70 dB SPL in 5 dB steps. Acoustic stimuli were delivered with a custom acoustic assembly consisting of two electrostatic drivers (EC-1, Tucker Davis Technologies) and DPOAEs were collected with a miniature ER 10B+ low noise mic system (Etymotic Research Inc.) designed to measure the level of the ‘cubic difference tone’ 2f1-f2.

For the contralateral-noise assay, ipsilateral DPOAEs were evoked with primaries (f1 = 12.73 kHz and f2 = 15,26 kHz) at tone level ranging from 30 to 70 dB SPL in 5 dB steps before, during and after contralateral noise application. Contralateral noise consisted of a continuous white noise delivered at 50 dB SPL directly to the contralateral ear canal with a super tweeter (Realistic^R^; frequency range 4-40kHz).

For the noise exposure assay, mice were exposed, awake, and unrestrained within a cage for 12h at night inside a small reverberant sound exposure box. The exposure stimulus was generated by a white-noise source at 120 dB SPL using a Marantz HD-493 speaker connected to an amplifier Marantz PM-493. Sound level exposure was measured within the cage using a Bruel & Kjaer 2238 Mediator Sound Level Meter.

### Confocal fluorescence imaging

#### Expression of P2X4mCherryIN in living (unfixed) cochlear tissue

Micro-dissection of the organ of Corti from freshly extracted cochlea from P6 to P60 mice was performed under binocular microscopy and in artificial perilymph at 4-8°C. The living freshly dissected organ of Corti was then placed and hold under dental-floss in a microscopy imaging-chamber to be observed under live confocal-fluorescence microscopy (excitation with a 543 nm Helium Neon Laser system (Melles Griot 05-LGR-193-381) coupled to a Nikon C2-confocal FN1-upright microscope and emission fluorescence analyzed at 552–617 nm) in order to monitor the direct expression of the fluorescent signal of mCherry protein fused to P2X4. LysoTracker probe Green DND-26 (Invitrogen, ThermoFisher Scientific, Catalog # L7526) was also used to simultaneously label lysosomes in live hair cells (incubation 15-25 min at 1-2 µM). The lysosomal dye was visualized under excitation with solid-state argon laser at 488 nm (laser 85-BCD-010-706, Melles Griot) and emission at 500-550 nM.

#### Carnoy’s fixation

In some experiments as indicated in the text, direct visualization of P2X4-mCherry was done after a 15 min incubation-perfusion of the dissected cochleae in Carnoy’ fixative solution. We found that Carnoy’s fixation allowed a better conservation of mCherry fluorescence signal as compared to paraformaldehyde (PFA) fixation. The cochleae were then rinsed with PBS and decalcified for 5 hours at RT with a PBS containing 5% EDTA. Hemi-section of the cochleae were then performed in order to observe in transversal sections the organ of Corti.

#### Immuno-detection of P2X4 in fixed cochlear tissue

Cochlear samples from P30-P60 mice were fixed by incubation in 4% PFA in phosphate-buffered saline (PBS), pH 7.4, at 4°C overnight and washed with cold phosphate buffered saline (PBS). They were then incubated overnight in PBS solution containing 10% EDTA pH 7.4 at 4°C for decalcification. The organ of Corti was then dissected and the tectorial membrane removed. P2X4mCherryIN expression was directly visualized by the endogenous fluorescence of mCherry protein or using an anti-RFP antibodies (Rabbit anti-RFP MBL #PM005). WT P2X4 receptors were labeled using an antibody recognizing the extracellular domain of native P2X4 (rat anti-P2X4 antibody Nodu-246, 1:200; obtained in collaboration with F. Koch-Nolte, Hamburg, Germany) (Bergman et al., 2019, Bertin et al, 2021; 2022) and the outer hair cells are revealed using phalloïdin (Invitrogen A22283, 1: 200) which marks the actin filaments of the stereocilia of the hair cells. After incubation with their respective fluorescent secondary antibodies (Anti-rat AlexaFluor568 Life technologies Catalog # A11006 or anti-rabbit AlexaFluor568 Life technologies Catalog # A-11036): the organs of Corti are mounted between slide and coverslip and analyzed by confocal imaging (Leica SP8, Bordeaux Imaging Center) for the expression and localization of the receptor of interest. To label the MOC efferent fibers contacting OHCs a specific rabbit monoclonal antibody against Choline Acetyl Transferase (ChAT) was used (Sigma, ZRB1012; 1:200) and a secondary donkey antibody conjugate with Alexa Fluor 488 (Molecular Probes, A-21206, 1:1000).

### Statistical analysis

Two-way analysis of variance (ANOVA) was used with the Šídák multiple comparison test to perform a statistical comparison under the statistical analysis software Graphpad Prism®. The independent two-sample *t*-test analysis tests was used for comparing two samples with normal distribution while the Wilcoxon/Mann-Whitney test was used as a non-parametric test for comparing two samples with non-normal distributions (OriginLab software). p<0.05 values were considered statistically significant.

## RESULTS

### P2X4 expression in OHCs of the auditory organ

The direct observation under confocal fluorescence microscopy of surface preparations of living organs of Corti, freshly micro-dissected from constitutive (P30-P60) P2X4mCherryIN mice, showed a strong intracellular endogenous fluorescence of mCherry fused to P2X4 in the three rows of outer hair cells (OHCs) but not in the inner hair cells (IHCs) and the supporting cells (Fig.1). A similar cellular fluorescence pattern was observed from the base (high frequencies) to the apex (low frequencies) of the cochlea (Fig.1A). By focusing up and down into the living surface preparations of the organ of Corti, from the stereocilia to the basal synaptic end of the OHCs, we noticed that the fluorescent P2X4mCherry signal was essentially restricted below the cuticular plate, an area known to be rich in lysosomes and called the Hensen’s body. Note that this intracellular organelle is a whorl of lamellar and tubulovesicular cisternal ER, associated with mitochondria, where a Ca^2+^-mobilizing second messenger system linked to ATP-P2 receptor has been shown to take place (Mammano et al., 2019). The use of the dye Lysotracker to label the lysosomes indicated a good colocalization with the P2X4mCherryIN signal in OHCs (Fig.2C-2D). A similar P2X4mCherryIN fluorescent signaling in OHCs was found in Myo15-Cre^+/−^:P2X4mCherryIN mice, when the expression of the Cre recombinase was driven under the control of a hair cell specific promotor (Dulon et al., 2018; Fig.1A-C). No P2X4mCherryIN expression was found in OHCs of Syn1-cre^+/−^:P2X4mCherryIN mice, when the promotor of the Cre recombinase was neuron specific (Fig.1F). Also, no fluorescent labeling was found in OHCs of WT mice (P2X4mCherryIN^fl/fl^ not expressing the cre recombinase) (Fig.1G). The specific expression of P2X4 in OHCs was confirmed by confocal immunofluorescence microscopy when using an antibody directed against P2X4 (anti-P2X4 Nodu-246) while no staining was revealed using the same antibody on constitutive P2X4KO mice (Fig.2A-2B). The intracellular labeling in the region of the Hensen’s body of the cylindrical OHCs was confirmed in sagittal sections of the cochlea from P6 to P60 P2X4mCherryIN mice (Fig.3A-3C). The expression of P2X4mCherryIN in OHCs was also found at the plasma membrane and in particular at the basal post-synaptic cholinergic pole of OHCs facing the MOC efferent fibers expressing ChAT (Fig.3D-3E). In P2X4KO mice, the absence of P2X4 did not modify the organization/distribution of the cholinergic efferent fibers/terminals contacting the OHCs (Fig.3F-G). Note that P2X4mCherryIN was transiently present in immature developing IHCs of pre-hearing P6 mice (Fig.3C) while being absent in adult post-hearing IHC from P60 mice (Fig.3A).

**Figure 1:**
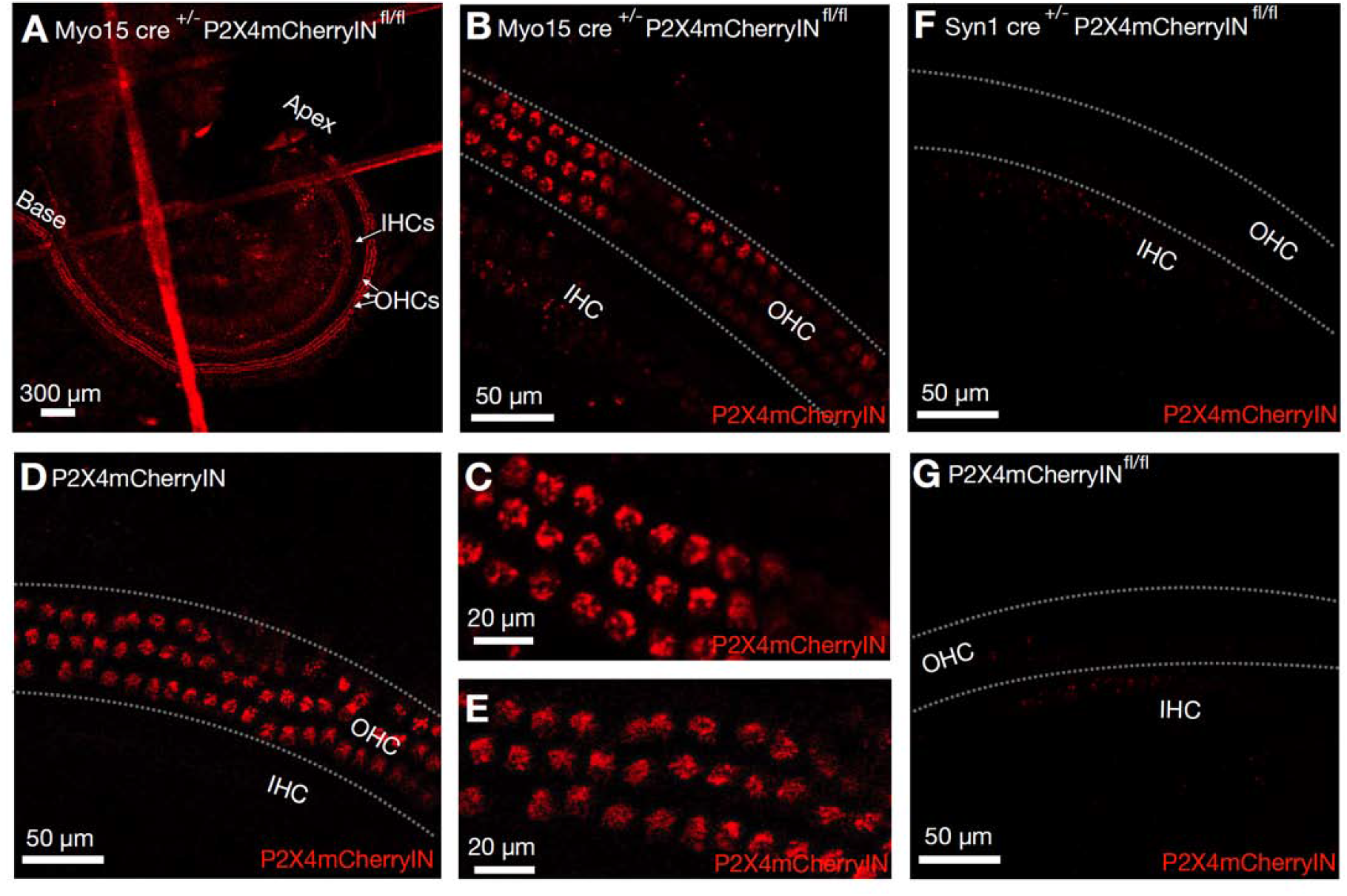
P2X4mCherry is selectively expressed in OHCs of living organ of Corti. Direct visualization with confocal laser microscopy of the endogenous fluorescence of P2X4mCherryIN in living organ of Corti explants freshly dissected from different transgenic mice (P60): (**A**,**B**,**C**) Myo15-cre^+/−^:P2X4mcherryIN^f/f^ mice showed specific expression of P2X4mCherry in OHCs but not in IHCs or supporting cell of the organ of Corti; (**D**,**E**) General (constitutive) CMV^+/−^ :P2X4mcherryIN mice showed similar expression of P2X4mCherry in OHCs; (**F**) Syn1-cre^+/−^ :P2X4mcherryIN^f/fl^ mice, with neuronal-specific expression of cre-recombinase, showed no labeling in the organ of Corti; (**G**) Control P2X4mcherryIN^fl/fl^ mice showed, in absence of cre expression, no fluorescent P2X4mCherry.

**Figure 2:**
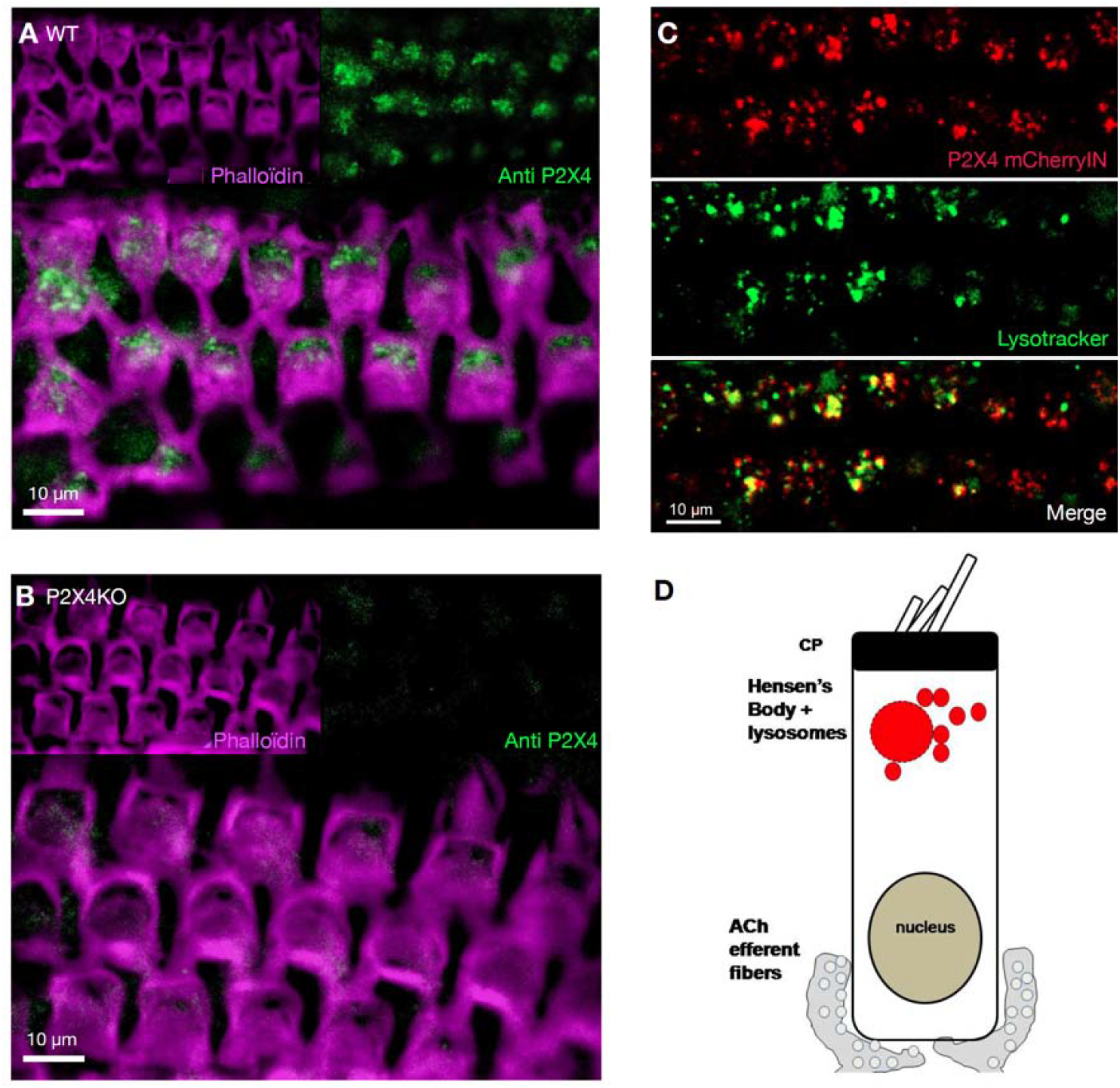
P2X4 is mainly localized in lysosomes in OHCs. (**A-B**) Confocal images of immunofluorescence labeling of P2X4 receptor using anti-P2X4 Nodu-246 antibody (green) and F-actin with phalloidin labeling (purple) of PFA-fixed organ of Corti surface preparations (P60), from wild-type (**A**) and P2X4KO mice (**B**). (**C**) Surface preparation of freshly dissected living organ of Corti of a general CMV^+/−^ :P2X4mcherryIN mouse (P60) showing co-localization of endogenous fluorescence of P2X4mCherryIN (red) and lysotracker-labeled lysosomes (green) in OHCs. (**D**) Schematic drawing of an OHC and its lysosomal complex expressing P2X4mCherry (red) below the cuticular pate (CP).

**Figure 3:**
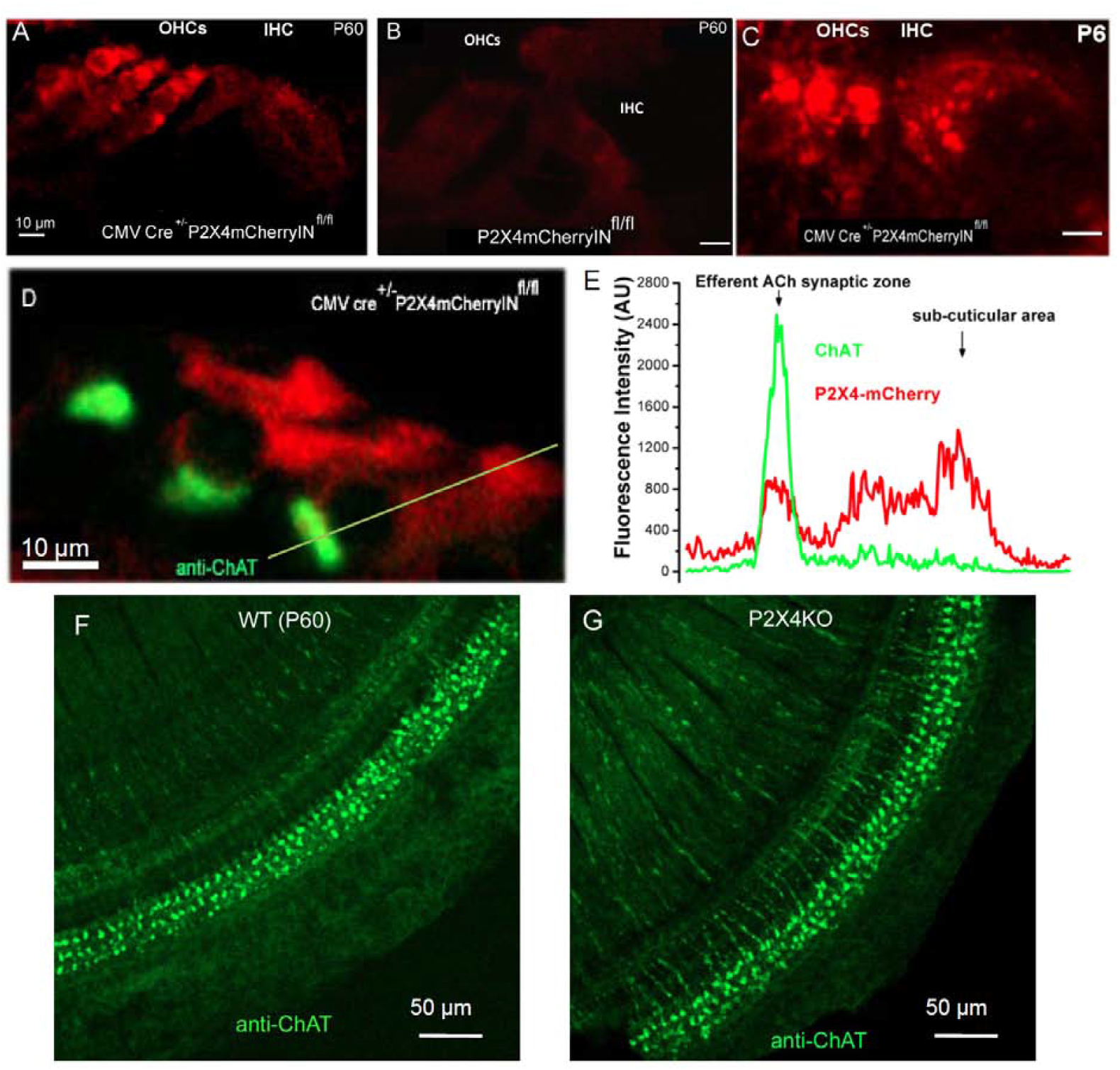
P2X4 in OHCs are facing the ChAT expressing MOC efferent terminals. (**A**) Confocal images of immunofluorescence of P2X4mCherryIN receptors using anti-RFP antobodies in PFA-fixed hemi-cochlea sections of general CMV Cre^+/−^ P2X4mcherryIN and in control P2X4mCherryIN^fl/fl^ mice (**B**). (**C**) Direct visualization of P2X4mCherry in a living (unfixed; freshly dissected) hemicochlea section from a P6 general CMV Cre^+/−^ P2X4mCherryIN mouse. (**D**) Confocal images of immunofluorescence of P2X4mCherryIN (red) and anti-ChAT antibodies (green) in a (Carnoy’s) fixed section of the organ of Corti of P60 general CMV Cre^+/−^ P2X4mCherryIN mouse. The fluorescence intensity profile (**E**) measured on the yellow line in **D** shows the co-localization of the cholinergic efferent nerve terminals and P2X4 at the base of the OHCs. (**F-G**) Confocal images of ChAT immunolabeling of PFA-fixed organs of Corti surface preparation from WT and P2X4KO mice showed similar efferent cholinergic terminals below OHCs.

### Constitutive and hair cell specific deletion of P2X4 in mice increased cochlear DPOAEs

To reveal the function of P2X4, we characterized the sensory and neural impact of P2X4 deletion in 60-day-old mice by measuring auditory evoked potentials (ABRs) and otoacoustic distortion products (DPOAEs). Surprisingly, click-ABR responses of constitutive P2X4KO mice (n=11) showed a significant potentiation with a decrease in latencies and thresholds as compared to control (P2X4mCherryIN^fl/fl^) mice (n = 11; Fig.4A-C). This hyperacusis like-effect was associated with a large increase in the amplitude of the DPOAEs (Fig.5A-B), which likely takes origin from an increased electromechanical activity of the OHCs. Similar increase in the amplitude of the DPOAEs was observed when deleting P2X4 selectively in hair cells of Myo15-cre^+/−^:P2X4KO^fl/fl^ mice (n = 7; Fig.4C), reinforcing the hypothesis of a specific role of P2X4 in OHC micromechanics. Of note, Syn1-cre^+/−^:P2X4KO^fl/fl^ mice (n = 13), in which P2X4 is selectively deleted in neurons, DPOAEs remained normal (Fig.4D). Also, in these latter mice, both the amplitude and latency of tone and click-evoked ABRs were comparable to control (P2X4mCherryIN^fl/fl^ and WT) mice (data not shown), suggesting that the deletion of P2X4 in cochlear and central neurons does not affect hearing.

**Figure 4:**
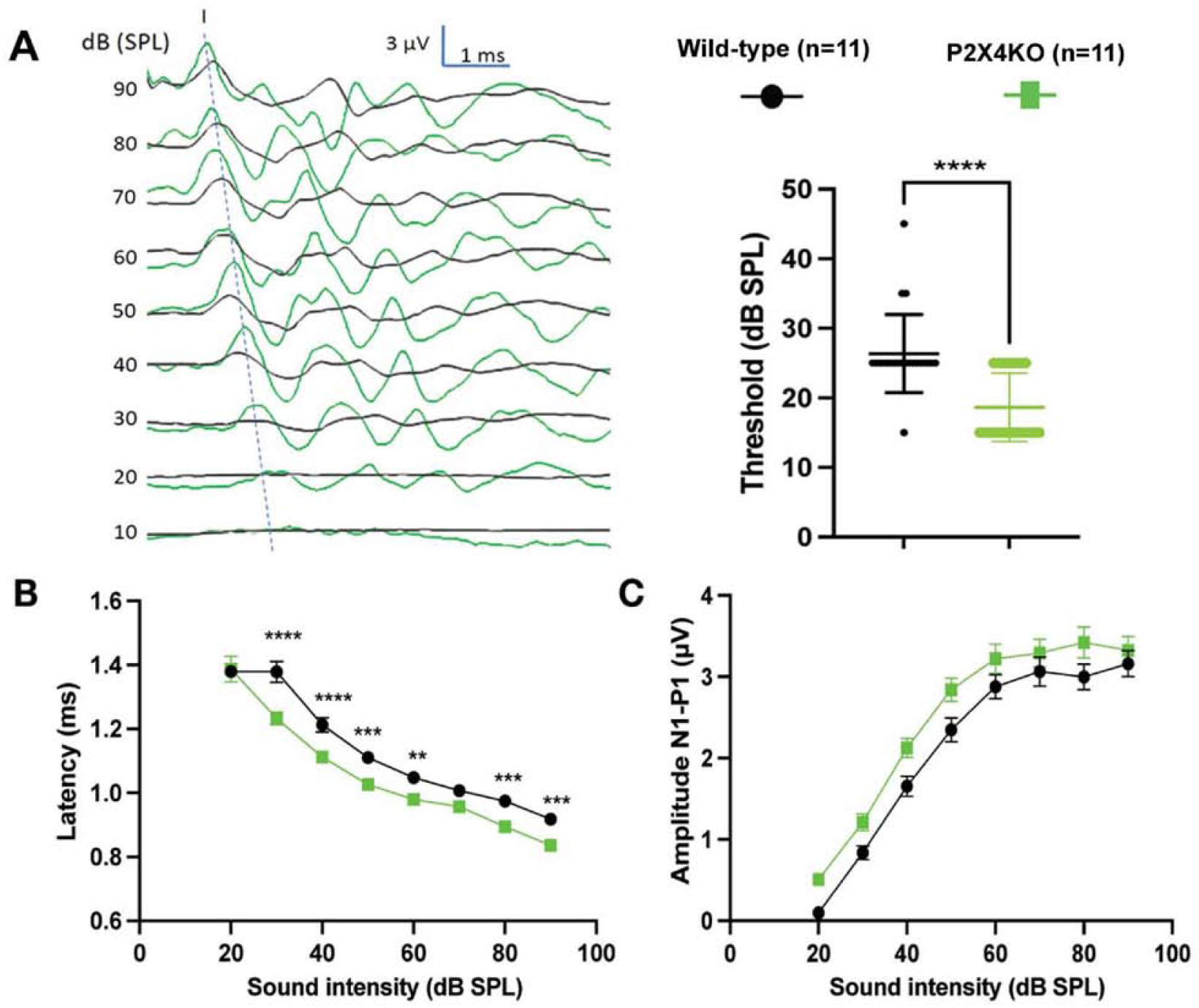
Comparative Auditory Brainstem Responses (ABRs) in WT and P2X4KO mice. (**A**) At left, examples of click-evoked ABR waves in a P60 wild-type mouse (black) and a P60 P2X4KO mouse (green). At right, mean comparative thresholds of click-evoked ABRs ((****, p<0.0001, two-sample *t*-test analysis). (**B**) Comparative latencies of wave-I as a function of sound intensity (**, p<0.01, ***, p<0.001, two-way ANOVA). (**C**) Comparative wave-I amplitudes as a function of sound intensity were not significantly different (two-way ANOVA).

**Figure 5:**
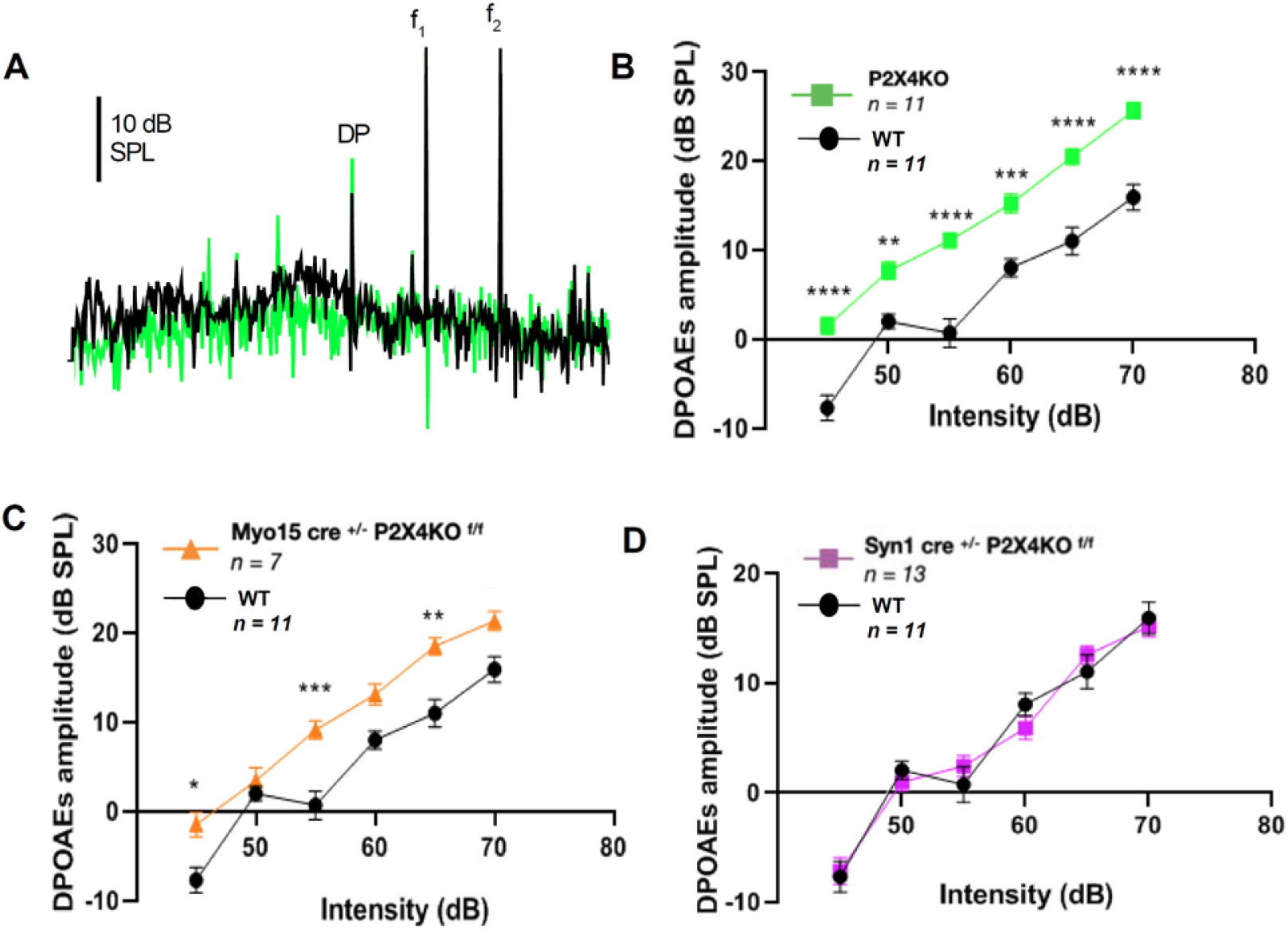
Distortion Product Otoacoustic Emissions (DPOAEs) generated by OHCs, are augmented in P60 general and OHC-specific P2X4KO mice. (**A**) Examples of DPOAEs (Distortion Product (DP) = 2f1-f2) recorded at f_2_ = 15.26 kHz (65 dB SPL) and f_1_= 12.73 kHz recorded in WT and mutant mice. (**B**) Comparative amplitude as a function of sound intensity of f_1_f_2_ primaries in constitutive P2X4KO mice and WT mice. (**C**) Conditional hair cell specific deletion of P2X4 lead to similar increase in DPOAEs. (**D**) DPOAEs were similar to WT in Syn1 cre^+/−^ P2X4KO^f/f^ mice, suggesting that the increased DPOAEs in B-C do not result from a neuronal defect but rather from OHCs. (**, p<0.01, ***, p<0.001, **** p<0.0001; two-way ANOVA).

### Contralateral noise suppression of DPOAEs is altered in P2X4KO mice

In the organ of Corti, OHCs receive efferent innervation from the medial olivocochlear (MOC) system, a group of cholinergic neurons originating in the superior olivary complex (Guinan, 2006). MOC neurons are activated by sound and they are known to adjust cochlear sensitivity and protect from acoustic overstimulation (Maison et al., 2013). This cholinergic MOC reflex can be assayed noninvasively via its suppressive effects on DPOAEs (Maison et al., 2012). To assess whether P2X4 deletion affected the MOC suppressive effects on cochlear OHC responses, DPOAE amplitudes were measured before, during and after the presentation of a continuous broadband noise (50 dB SPL) to the contralateral ear in wild-type and P2X4KO mice (Fig.6). To evoke ipsilateral DPOAEs, primary tones (f_1_= 12.73 kHz and f_2_=15.26 kHz) were presented continuously at a sound pressure of 55 dB SPL (Fig.6A-B) or 65 dB SPL (Fig.6C). The contralateral noise suppression of the DPOAEs was much less effective in P2X4KO mice as compared to WT mice, (n = 5 for each genotype; p<0.001; Fig. 6A-C). To confirm this diminished contralateral suppression of DPOAEs in P2X4KO mice, we performed another set of contralateral masking experiments with different P60 WT and mutant mice in which we varied the intensities of the primaries from 30 to 70 dB SPL. At low primary sound levels ranging from 40-55 dB SPL, while DPOAEs in WT mice were completely suppressed by a 50 dB contralateral noise (n = 9), significant DPOAEs could still be recorded in P2X4KO mice (n = 10; Fig.6D). These results suggested that P2X4 are essential for an efficient efferent MOC suppression of DPOAEs.

**Figure 6:**
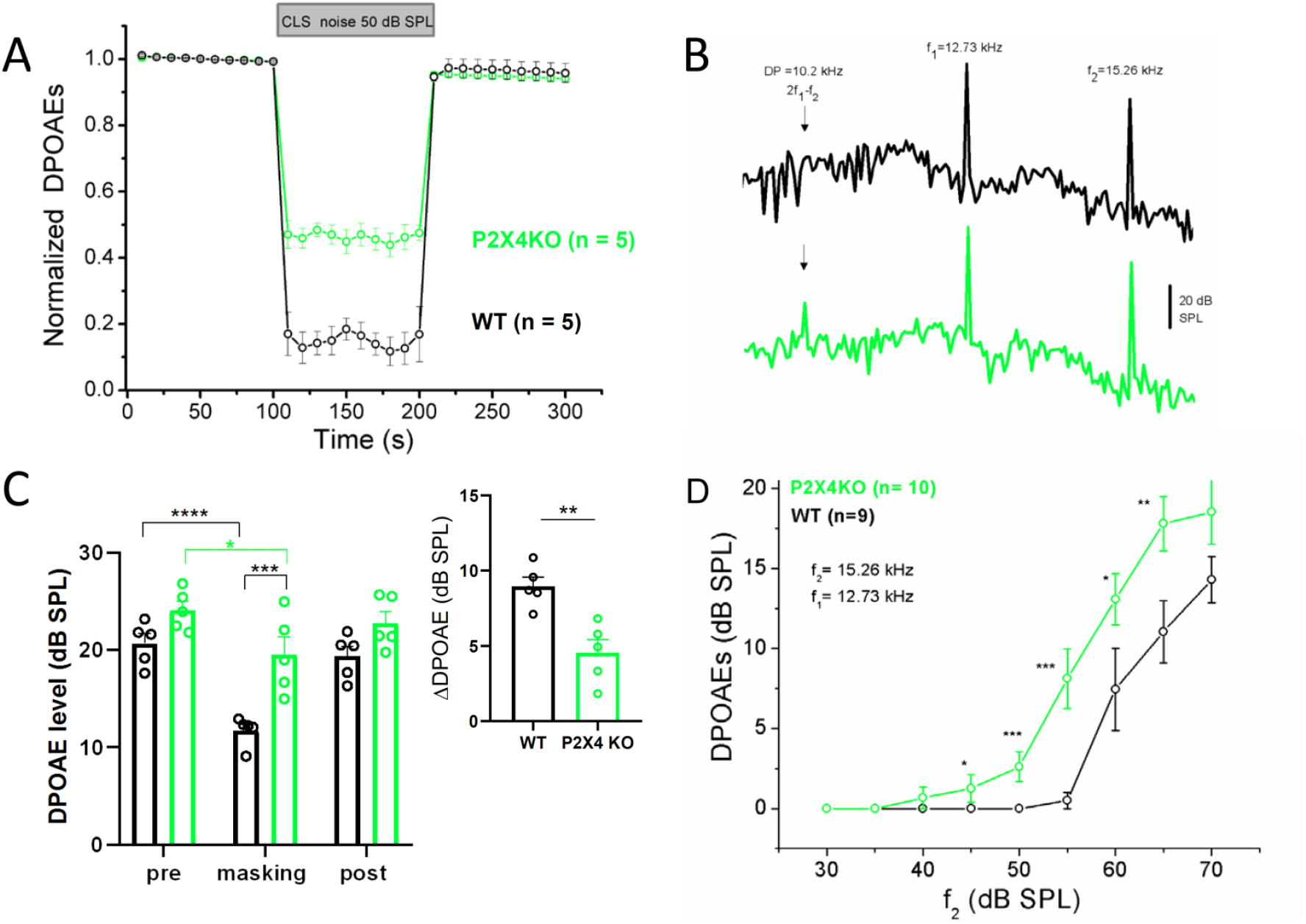
The efficiency of contralateral-noise suppression of ipsilateral DPOAEs was reduced in P2X4 KO mice. (**A**) Comparative reduction of DPOAEs in WT (black) and mutant mice (green) when transiently applying a low-intensity white noise (50 dB SPL) to the contralateral ear. Ispsilateral DPOAEs were evoked with primaries intensity set at 55 dB SPL (f_1_= 12.73 kHz and f_2_=15.26 kHz). DPOAEs were continuously measured 10 times (one per s) before, during and after the application of the contralateral noise. (**B**) Typical examples of DPOAEs (from A) during contralateral noise application. Note the absence of DP in the WT (black) as compared to the P2X4 KO (green). (**C**) At left, mean DPOAEs levels before (pre), during masking and after (recovery); same mice and conditions as in A but DPOAEs evoked with primaries at 65 dB SPL; n = 5 for each genotype; one-way ANOVA). At upper right, graph shows comparative suppression level (pre-level DPOAEs minus masking levels) for each genotype. Note the significant lower suppression in mice lacking P2X4 (two-sample *t*-test analysis). (**D**) In a different group of mice (n = 9 WT and n = 10 P2X4KO), comparative suppression of DPOAEs as a function of ipsilateral primaries intensity during a constant application of a contralateral white noise of 50 dB SPL The suppression by contralateral noise was reduced at all intensities in P2X4 KO mice (two-way ANOVA; * p<0.05; ** P<0.01; *** P>0.001).

### Faster recovery of DPOAEs after noise exposure in P2X4KO mice

We then evaluated the effects of P2X4 deletion on the vulnerability of the hearing function to acoustic noise injury. WT and P2X4KO mice (P60) were continuously exposed to a white noise at 120 dB SPL for 12h and allowed to recover up to 14 days. DPOAE and click ABR amplitudes were assessed before, one hour after noise exposure and every 24 h or 36 h up to 14 days (Fig. 7). Both WT and mutant animals showed no DPOAEs when tested immediately after noise exposure and 24 hours later, indicating significant damage in the OHC’s function. Unexpectedly, mutant mice lacking P2X4 showed a faster recovery of their DPOAEs (temporary threshold shift (TTS) with half-time recovery ~ 3 days) as compared to wild-type (TTS half-time recovery ~ 6 days) (Fig. 7A). Both WT and P2X4KO mice recovered to a similar sustained DPOAEs level after 8 days, this level corresponding to about 75% of their pre-exposed amplitudes after 14 days, leaving a permanent threshold shift (PTS). In the same mice, the impact of noise exposure was also measured on click-evoked ABR wave I amplitude. ABR responses in mutant and WT mice showed a similar reduction and recovery pattern when tested with a click at 85 dB SPL (Fig. 7B). It is to be noted that this high sound intensity by pass the OHC amplification process and directly probed the function of the IHC ribbon synapses. These results suggested that the absence of P2X4, contrary to the DPOAEs, did not influence the time recovery of the IHC’s afferent synapses after noise exposure.

**Figure 7:**
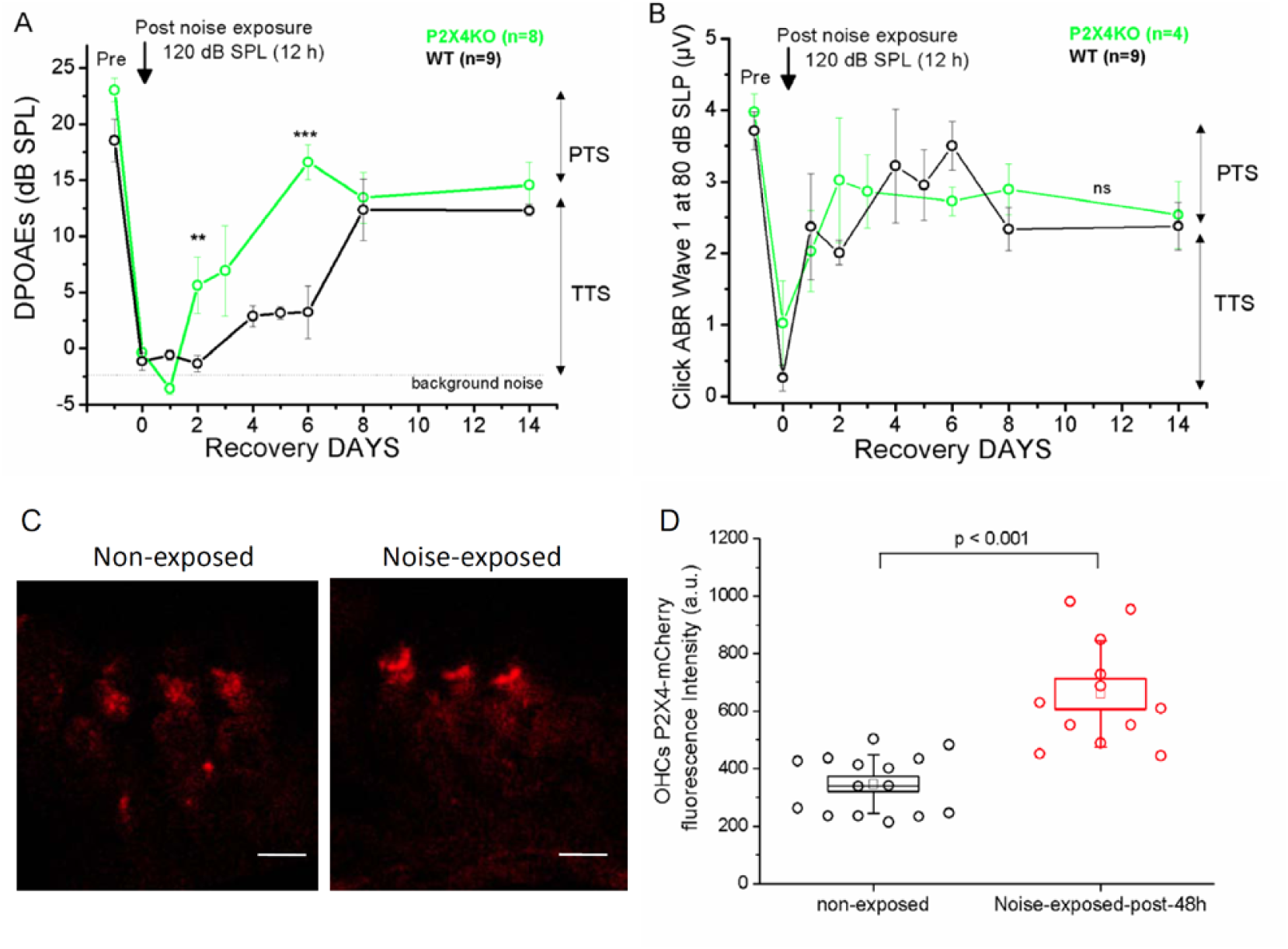
Recovery kinetics of auditory function in P2X4 KO and WT mice (P60) following noise exposure (120 dB SPL for 12 hours). (**A**) Faster recovery of DPOAEs evoked in P2X4KO mice after noise trauma as compared to WT mice (** p<0.01, *** p< 0.001 with two-sample t-test comparison at 2 and 6 days post-noise; n=8 for WT and n=9 for P2X4KO mice): The evolution of the amplitude of the DPOAEs before and after noise are here shown with primaries set at 65 dB SPL. (**B**) Click ABR recordings in P2X4KO mice showed a similar pattern in the recovery time as compared to WT mice. The evolution of the ABR amplitudes are here shown at 80 dB SPL. (**C-D**) Fluorescence images were obtained from sagittal sections of cochleae (Carnoy’s fixation) of general P2X4mCherryIN mice (P60) in control condition and after noise-exposure(white noise 120 dB SPL, 12h). (**D**) The endogenous fluorescence of mCherry protein fused to P2X4 in OHCs revealed an increased expression of P2X4mCherryIN after noise-exposure. The intensity of fluorescence was measured in the subcuticular region (square 5 µm^2^) of the OHCs. The fluorescence intensity in individual OHCs was compared between non-exposed (black) and noise-exposed (red) mice, (12–15 cells analyzed per condition from three mice in each group; two-sample t-test comparison with *** p<0.001). Temporary Threshold Shift (TTS); Permanent Threshold Shift (PTS).

### Overexpression of P2X4 in OHCs after noise exposure

To determine whether noise exposure could affect the expression of P2X4 in OHCs, we used general P2X4mCherryIN (P60) mice where the expression of mCherry-tagged P2X4 was used as a direct fluorescent reporter of P2X4 expression. We found that noise exposure (white noise, 12 h at 120 dB SPL) significantly increased the expression of P2X4mCherryIN in OHCs after 48 h (Fig. 7C-D). This overexpression of P2X4 in OHCs after noise exposure in WT mice may produce a sustained reduction of DPOAEs and explain the slower recovery of DPOAEs after noise exposure in WT mice as compared to P2X4KO mice.

## DISCUSSION

### Specific expression of P2X4 in OHCs

Intracochlear purinergic receptors, both ionotropic (P2X) and metabotropic (P2Y), have long been recognized as important mediators in regulating several aspects of hearing function from development to adult stages (Ito and Dulon, 2010; Jovanovic and Milenkovic, 2020; Vlajkovic and Thorne, 2022; Babola et al., 2021). Current studies indicate direct modulation of both IHCs and OHCs and their respective spiral ganglion neurons through yet not fully revealed combinations of P2 receptors. In hair cells, P2X2 (Järlebark et al., 2000) and P2X3 (Huang et al., 2003; Wang et al., 2020) are mostly expressed in the developing cochlea, while P2X4 have been described in the adult OHCs (Szücs et al., 2004; Ziyin et al., 2024) and spiral ligament capillaries (Wu et al., 2011) but their role remain unknown. Although we found a transient expression of P2X4 in immature P6 IHCs, spontaneous spiking activity of developing IHCs is not affected in mice lacking P2X4 (Sendin et al., 2014). Our present study showed a strong and specific expression of P2X4 receptors in both immature and adult mouse OHCs, by directly visualizing the endogenous fluorescence of mCherry protein fused to P2X4 in living OHCs from freshly dissected surface preparation of organs of Corti of general and OHC-specific P2X4mCherryIN Knockin mice or indirectly by using confocal immunohistochemistry with specific anti-P2X4 antibodies in PFA-fixed cochlear tissue of WT mice. Previous studies on P2X4mCherryIN mice have shown that under baseline conditions, fluorescence from P2X4mCherryIN is undetectable in the brain and spinal cord but is directly visible in isolated peritoneal macrophages highly expressing P2X4 (Bertin et al., 2021; Bertin et al., 2022; Gilabert et al. 2023). Notably, endogenous P2X4mCherry fluorescence has been observed in brain microglia following LPS-induced inflammation, a condition that triggers microglial activation and de novo *P2X4* expression (Bertin et al., 2021). In our present study, P2X4 were found mainly localized intracellularly, associated with lysosomes, in the Hensen’s body region of the OHCs. This localization of P2X4 in OHCs fit well with their well documented specific subcellular distribution, that is predominantly intracellular, within endolysosomal compartments of many cell types (<u>Qureshi et al., 2007</u>; Murrell-Lagnado 2018). Even in P2X4mCherryIN mice where internalization is defective and surface expression is expected to increase (Bertin et al., 2021), P2X4mCherryIN fluorescence remains predominantly intracellular in OHCs. Lysosomes are considered as potent important Ca^2+^ stores, potentially releasing Ca^2+^ in the cytosol upon the activation of endolysosomal P2X4 via luminal ATP and pH alkalinization (Huan g et al., 2014; Cao et al., 2015; Murrell-Lagnado 2018). Endolysosomal P2X4 receptors can be activated downstream of plasma membrane Ca^2+^ permeable P2X7 receptor in rat kidney (NRK) fibroblasts and the increase in near-lysosome cytosolic [Ca^2+^] is exaggerated via release through P2X4 (Tan et al., 2021). Since we found that P2X4 are also present at the OHC plasma membrane, notably facing the ChAT MOC efferent terminals, where Ca^2+^ permeable nAChRs are also expressed (Blanchet et al., 1996; Vyas et al., 2020), the Ca^2+^ influx through P2X4 or/and nAChRs could potentially be amplified by the intracellular P2X4 endolysosomal compartments (see hypothetical representation in Fig. 8).

**Figure 8:**
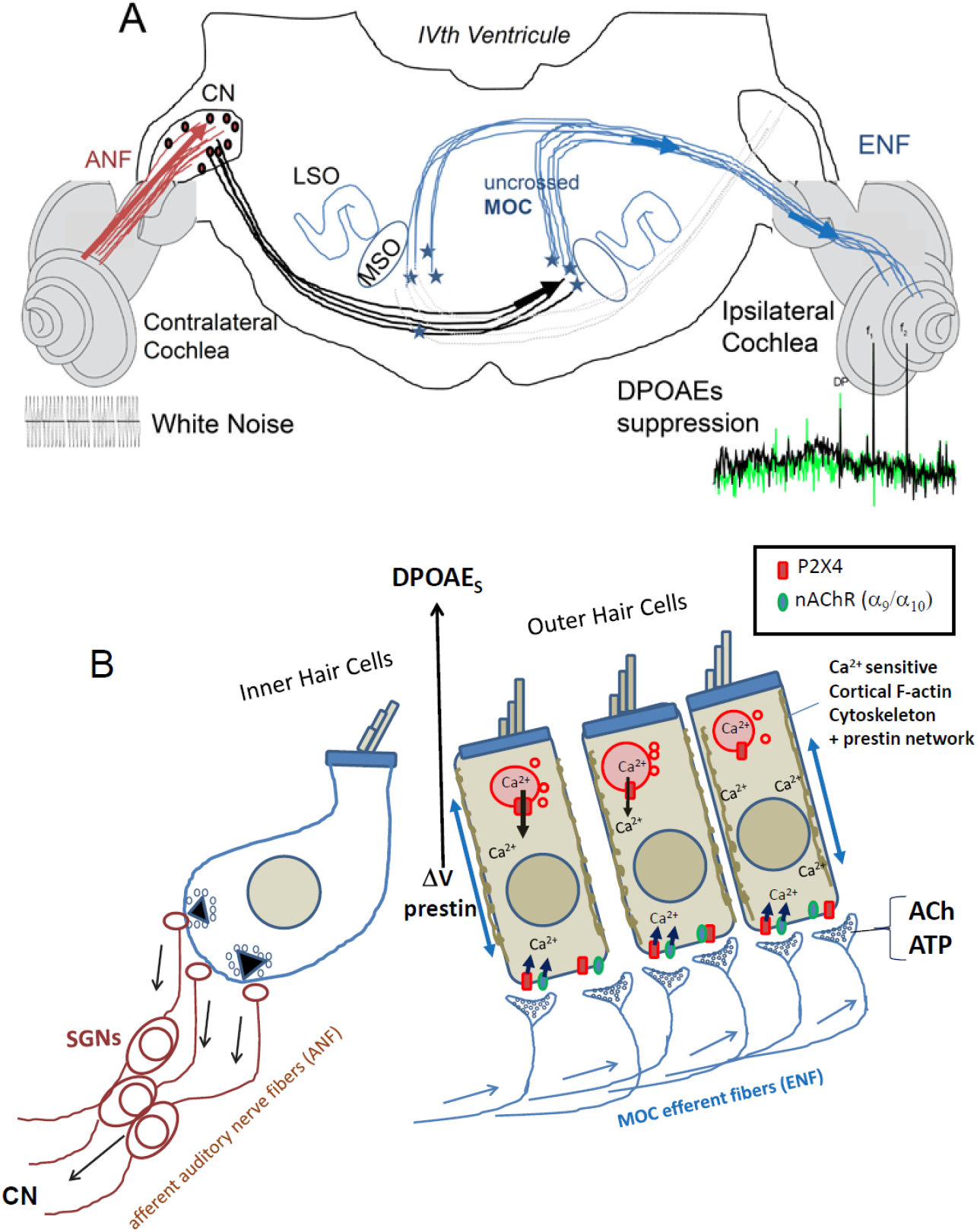
Hypothetical efferent mechanisms regulating cochlear OHCs’ micromechanics and DPOAEs via P2X4 activation. (**A**) Schematic drawing of the medial olivocochlear (MOC) reflex pathways for sound evoked-controlateral suppression to the DPOAEs of the ipsilateral cochlea. Cochlear nucleus (CN) neurons, receiving inputs from the auditory nerve fibers (ANF), project to MOC neurons on the opposite side of the brain. These MOC neurons project inhibitory cholinergic efferent nerve fibers (ENF) to OHCs of the ipsilateral cochlea. LSO, lateral superior olive; MSO medial superior olive. (**B**) Schematic drawing of the organ of Corti located inside the cochlea shows the arrival of the inhibitory MOC efferent fibers below the OHCs. Both ACh and ATP could be released by these efferent fibers, in turn activating Ca^2+^ permeable P2X4 (red) and nAChRs (green) co-localized at the OHCs’ postsynaptic membrane. This Ca^2+^ influx will then activate nearby SK and BK channels (not shown) leading to OHC hyperpolarization and the inhibition of the voltage-driven prestin-electromotility and DPOAEs. The Ca^2+^ response could in addition be amplified by intracellular lysosomal P2X4 and leads to a change in stiffness of the F-actin cortical network of OHCs, reducing then the prestin-electromotility.

#### Constitutive and conditional P2X4 KO mice display augmented DPOAEs

This specific P2X4 expression in OHCs prompted us to examine the effects of their deletion on the DPOAEs in P2X4KO mice. Cochlear DPOAEs arise from the electromechanical activity of the OHCs, which amplify basilar membrane vibrations enhancing hearing sensitivity and frequency selectivity. Remarkably, we found that the DPOAEs in constitutive and OHC-specific P2X4KO mice were significantly augmented, suggesting that the expression of P2X4 receptors may down regulate electromotility in OHCs, a voltage-dependent mechanical process known to be regulated by Ca^2+^ (Dulon et al., 1990; Frolenkov et al., 2003; Sinha and Frolenkov, 2024). Recently, knock out of P2X7 receptors has also been shown to increase hearing sensitivity by increasing OHC electromotility (Liang et al., 2024). In this latter study, however, P2X7 receptors were found to be extensively expressed in the presynaptic efferent MOC terminals contacting the OHCs through which they may regulate their mechanical activity by controlling ACh and ATP release.

#### Impairment of contralateral-noise suppression of DPOAEs in P2X4KO mice

Since DPOAEs are known to be down-regulated by the activation of the MOC efferent system descending from the medial superior olivary nucleus (Galambos, 1956; Avan et al., 1996; Guinan 2006; Maison et al., 2003; see schematic representation in Fig.8A), we explored whether contralateral-noise suppression of DPOAEs was affected in P2X4KO mice. Note that one major component of this inhibitory efferent system is cholinergic and mediated by the α9/α10 nicotinic acetylcholine receptor (nAChR) complex expressed by OHCs (Vetter et al., 1999; Maison et al., 2002). This unique nAChR complex is functionally coupled to calcium-activated potassium (K_Ca_) channels, in particular SK2 channels, at the basal synaptic pole of the OHCs (Dulon et al; 1998; Kong et al., 2008; Oliver et al., 2000; Murthy et al., 2010). Calcium entry through the nAChR complex triggers K^+^ efflux resulting in OHC hyperpolarization (Housley and Ashmore, 1991; Fuchs and Murrow, 1992; Blanchet et al., 1996). This process regulates the voltage-dependent prestin-based membrane motors which are fundamental to the generation of DPOAEs. Mice lacking α9 nAChR fail to show suppression of DPOAEs (Vetter et al., 1999) and mutation in the α9/α10 nAChR with increased duration of channel gating (Taranda et al., 2009) greatly increases the time course of efferent MOC suppression (Wedemeyer et al., 2018). In our present study, we showed that contralateral-noise suppression of DPOAEs evoked in mice lacking P2X4 is also largely reduced, suggesting that a Ca^2+^ dependent activation of K_Ca_ channels in OHCs may also occur via Ca^2+^ entry and/or intracellular release through P2X4 (Fig.8B). This P2X4-dependent process acting on DPOAEs is likely to occur at the level of the OHCs, as it was observed exclusively in conditional P2X4KO mice with hair cell-specific Myo15-cre deletion and not in those with neuronal Syn1-cre deletion (Fig. 5). Also, the effects on DPOAEs of P2X4KO mice were unlikely due to a loss of MOC cholinergic efferent synapses since the number of ChAT efferent boutons below the OHCs was similar in WT and mutant mice (Fig. 3F-G). Co-release of ATP and ACh, acting at postsynaptic P2X-containing and nACh receptors has been suggested to play a dominant role in brain (Rodrigues et al., 2006) and carotid body chemoreceptors (Zhang et al., 2000). We propose that a similar co-release mechanism, involving P2X4 and nAChRs, takes place at the MOC efferent terminals contacting the OHCs (Fig. 8B).

#### Faster recovery of DPOAEs after noise trauma in absence of P2X4

A role in protection from acoustic injury has been proposed for the MOC efferent system (Reiter and Liberman, 1995). One major component of this protection is mediated by the α9/α10 nAChR complex since overexpression of α9 nAChRs in OHCs significantly reduces acoustic injury from noise exposure causing either temporary or permanent damage (Maison et al., 2013).

Recently, P2X7 have also been suggested to be important for hearing sensitivity and noise protection, likely through its expression at type II afferent cochlear neurons (Liang et al., 2024). These type II neurons, which innervate OHCs, provide inputs to the central MOC efferent system and are thought to play the role of nociceptive fibers reporting cochlear overstimulation and damage (Liu et al., 2015). In our study, P2X4KO mice did not exhibit significantly increased temporary (TTS) or permanent (PTS) auditory damage after noise exposure. On the contrary the P2X4KO mice showed faster time recovery of their DPOAEs after noise injury. The slower recovery time observed in WT mice after noise exposure could be attributed to P2X4 overexpression, a process recently reported in the cochlea using real-time quantitative PCR detection following noise trauma (Shi et al., 2024) and in OHCs (our data, Fig. 7C-D). An overexpression of P2X4 after noise exposure in WT OHCs may sustain a significant attenuation of DPAOEs for several days, providing protection against further noise-induced damage. In addition, since P2X receptors are known to physically and functionally interact with other ionotropic receptor-channels, notably through cross-inhibition of nAChRs, (Boué-Grabot et al., 2003, 2004; Rodrigues et al., 2016; Khakh et al. 2000), the deletion of P2X4 in OHCs could lead to up-regulation of nAChRs activity. This, in turn, may enhance MOC efferent protection against noise-induced injury.

## Acknowledgements

This work was in part financed by the Fondation pour l’Audition (FPA grant number: n° FPA IDA09 to DD), Agence Nationale pour la Recherche (ANR) grants (ANR 20-CE14-0016 and ANR-20-CE17-0013 to EBG). This work was also supported by CNRS, INSERM, University of Bordeaux, ‘Investments for the Future’ programs of the University of Bordeaux’s IdEx program (GPR BRAIN_2030) and by the COST Action CA21130 ‘P2X receptors as a therapeutic opportunity (PRESTO)’.

## Notes

**Conflict of Interest:** Authors report no conflict of interest.

**Funding sources:** This work was in large part financed by the Fondation pour l’Audition (FPA grant number: n° FPA IDA09 to DD), Agence Nationale pour la Recherche (ANR) grants (ANR-20-CE14-0016 and ANR-20-CE17-0013 to EBG). This work was also supported by CNRS, INSERM, University of Bordeaux, “Investments for the Future” programs of the University of Bordeaux’s IdEx program (GPR BRAIN_2030) and by the COST Action CA21130 “P2X receptors as a therapeutic opportunity (PRESTO)”.

### Competing Interest Statement

The authors have declared no competing interest.

